# An algorithm and application to efficiently analyze DNA fiber data

**DOI:** 10.1101/2021.12.29.474465

**Authors:** Teodor Kirilov, Anastas Gospodinov, Kiril Kirilov

## Abstract

The duplication of genetic information (DNA replication) is central to life. Numerous control mechanisms ensure the exact course of the process during each cell division. Disturbances of DNA replication have severe consequences for the affected cell, and current models link them to cancer development. One of the most accurate methods for studying DNA replication is labeling newly synthesized DNA molecules with halogenated nucleotides, followed by immunofluorescence and microscopy detection, known as DNA fiber labeling. The method allows the registration of the activity of single replication complexes by measuring the length of the “trace” left by each of them. The major difficulty of the method is the labor-intensive analysis, which requires measuring the lengths of a large number of labeled fragments. Recently, the interest in this kind of image analysis has grown rapidly. In this manuscript, we provide a detailed description of an algorithm and a lightweight Java application to automatically analyze single DNA molecule images we call “DNA size finder”. DNA size finder significantly simplified the analysis of the experimental data while increasing reliability by the standardized measurement of a greater number of DNA molecules. It is freely available and does not require any paid platforms or services to be used. We hope that the application will facilitate both the study of DNA replication control and the effects of various compounds used in human activity on the process of DNA replication.

## 1. Introduction

In the S-phase of the cell cycle, replication in eukaryotes starts from multiple sites called origins of replication and runs in parallel. The process is under strict control to ensure that every part of the genome is replicated only once in a cell cycle, and no parts remain unreplicated.

The origins of replication are set during the G1 phase when pre-replication complexes are assembled on multiple sites along the genome, in a process known as licensing. At the transition from G1 to S phases of the cell cycle, replication is triggered by the cyclin-dependent kinase (CDK) and Dbf4-dependent kinase (DDK), which activate the core replicative helicase – the MCM2-7 proteins, leading to the recruitment of additional factors and the formation of the active helicase – the CMG complex (Gambus, Jones et al. 2006), which unwinds DNA double helix during replication. The unwinding of DNA at the origin by the CMG complex allows the replisome assembly, including the DNA synthesizing polymerases and the factors coordinating the various activities in the molecular machine that carries out DNA replication. After initiation, replication forks then travel bidirectionally outwards from the origin (Gambus, Jones et al. 2006, Labib 2010).

However, the movement of replication machines along the DNA molecule is not unimpeded. Many intrinsic factors (e.g., specific DNA sequences and proteins that are tightly bound to DNA) as well as environmental factors (various chemical compounds) slow down or stop replication forks, causing a state known as “replication stress”.

A consequence of replication stress is the accumulation of DNA damage (Hills and Diffley 2014) as the unwound DNA at the sites of stopped replication complexes is more susceptible to being cut; specific proteins target and break down stopped replication forks; non-replicated portions of chromosomes are severed during mitosis. Damage caused by replication stress is considered to drive carcinogenesis, being the generator of genome instability required for the acquisition of the properties making a cell cancerous (Gorgoulis, Vassiliou et al. 2005, Macheret and Halazonetis 2015). On the other hand, excessive replication stress would not allow the completion of the S-phase, and the cell would die. This suggests that the rapidly dividing cancer cells (and therefore having higher levels of replication stress) would be more susceptible to replication stress-inducing compounds and thus could be selectively destroyed (Lecona and Fernandez-Capetillo 2014).

These determine the importance of studying the replication fork dynamics and the factors that affect it. The most direct method to do so is the DNA fiber labeling, a technique in which cells are fed halogenated thymidine analogs (using an appropriate labeling scheme (Quinet, Carvajal-Maldonado et al. 2017), which get incorporated into the newly synthesized DNA. Following labeling, cells are lysed, and their DNA is stretched on a glass surface. Then, newly synthesized DNA can be stained using antibodies that bind specifically to the incorporated thymidine analogs and the length of labeled fragments measured.

**Fig. 1.**
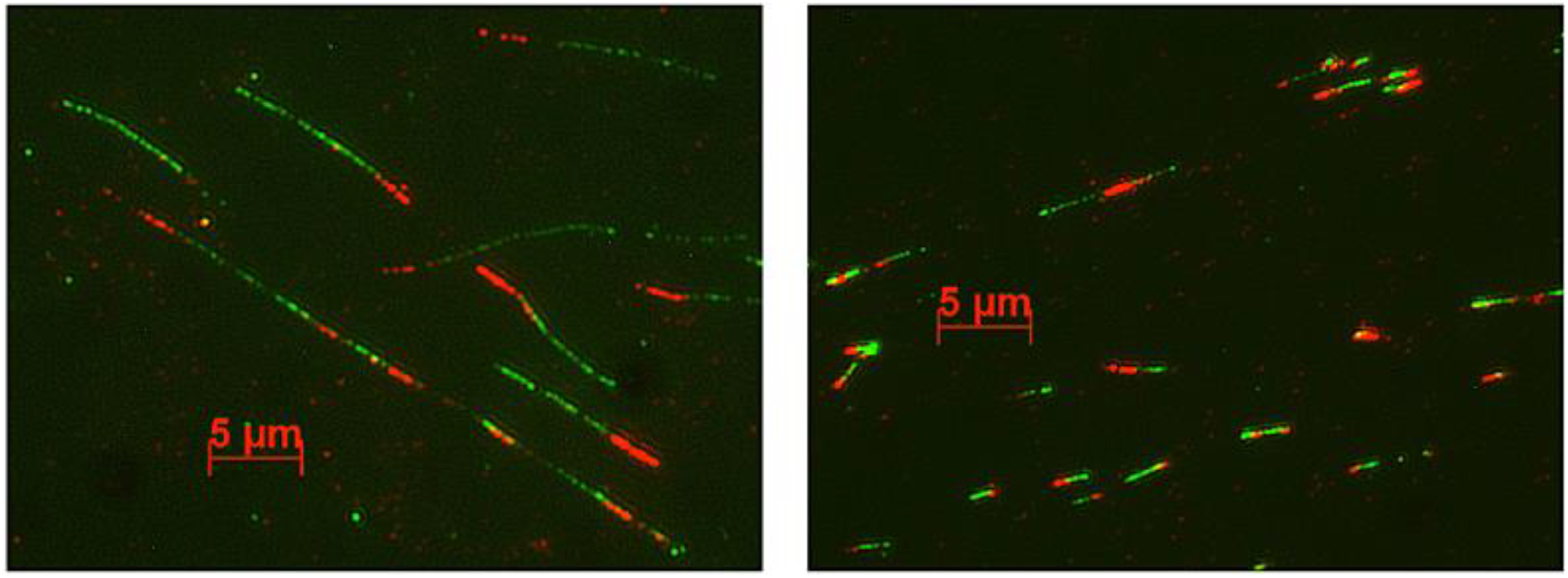
Typical images of nascent DNA molecules. The left image shows newly synthesized DNA from normally growing cells, and the right one shows nascent DNA molecules from cells experiencing replication stress (and therefore are shorter).

The main difficulty of this method is the complexity of image analysis -to obtain reliable data, it is necessary to measure many newly synthesized DNA fragments, which is time-consuming and requires highly trained researchers. In addition to being laborious, manual counting tends to introduce biases. Therefore, to increase the quality of fiber labeling data, it is necessary to increase the number of analyzed images to improve statistical reliability and use strictly set evaluation parameters.

To address the problems associated with the analysis of DNA fiber images, we developed “DNA size finder” - an application for image analysis written from scratch in Java, implementing our own algorithms to automate the image analysis. Thus, we provide a cross-platform, lightweight, easy to set up, easy to use, and completely free software solution to analyze DNA fiber labeling data. Furthermore, the users can opt for either a completely automated analysis or, by being presented with a detailed visual representation, can manually correct what the software “sees”.

## 2. Description of methods and algorithms

### 2.1. Software description

The entire application was written in Java (Version 14.0.2 64-bit) (Arnold, Gosling et al. 2000). The application and the algorithms were developed on Eclipse IDE for Java Developers (Version: 2020-06). Eclipse IDE for Java Developers is a free and open-source platform for the development of Java applications. The algorithms used are based on well-known methods from the image processing and graph theory, but unlike other proposed solutions of the same problem, we provide complete control over the analysis via many different parameters that the user can adjust.

### 2.2. Loading and initial processing of the images

After the image is loaded, it is represented as a matrix of pixels, each containing the information about its color in RGB format. The following notations are made:

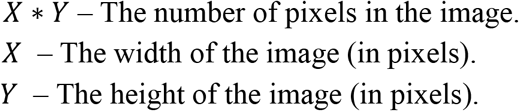

Due to variations in the color of original images obtained from a microscope, RGB binary thresholding is applied to analyze objects in color ranges, defined by minimum and maximum values set in the user profile. The complexity of the RGB binary thresholding is:

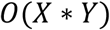

Since the reading of the input has the same complexity, the complexity of this thresholding process will not be included in the big O notation of the complexity of algorithms that use it a constant number of times.

### 2.3. Initial processing and finding objects of one color (classical approach)

The initial algorithm (which we also call “the classical approach”) will be described here, that we have developed for isolating and measuring single-color objects. This simple algorithm was the first that we created, and it was used as a basis for the development of more complicated algorithms that we use for analysis of the images.

Multiple artifacts are usually present in raw experimental data. To exclude artifacts, a “minimum object size” parameter is defined, through which they can be filtered. In addition, fibers in the image often appear split, although they belong to the same fiber. This necessitates the use of a second parameter, “Maximum Jump Distance”, based on which visually split objects are unified.

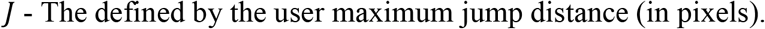

To achieve successful results, *J* needs to be kept low - between 5 and 15 pixels (typically corresponding to 0.5 micrometers). The original image is processed with the selected RGB binary thresholding settings. The image to be processed is presented as a non-oriented graph of type grid, in which every pixel represents a node. Edges exist between every two pixels if the Euclidean distance between those pixels (measured in pixels) is smaller or equal to the defined by the user *J*. The following notations are made:

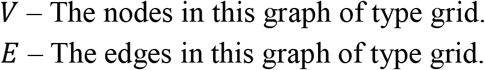

For every pixel, there is a node, so:

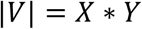

Every node creates edges to every node around in a circle with a radius *J*, so the number of edges coming from a node can be denoted with *O(J*^*2*^). For the number of edges of the entire graph can be made the approximation:

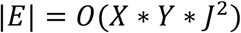

After the image is represented in this manner, the problem can be reduced to finding connected components in the resulting graph. This is processed through several breadths first searches, which are limited to traversing only the nodes that have passed the thresholding. Breadth-first search is preferred to depth-first search in order to avoid the use of recursive functions.

For each connected component found, the following are calculated:

I. First, its area is calculated by the number of pixels it is composed of. The connected components whose area is smaller than the minimum size specified by the user are removed.
II. The minimum and maximum values of the x and y coordinates of all of its pixels are calculated. Those values are used to define a rectangle that wraps the connected component, and its diagonal is used as the object’s length.

The connected components of the graph are found by using breadth-first search; hence the complexity of the algorithm for finding the objects is:

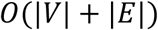

*V* and *E* can be replaced in this formula:

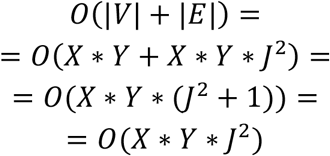

The complexity of this algorithm for finding objects of one color in the image by the classical approach is:

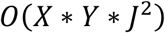

This complexity is both valid in the worst and best case scenarios since in this algorithm every node and edge in the grid will always be traversed.

### 2.4. Skeletonization-based approach for finding objects of one color

#### 2.4.1. An overview of the second algorithm for finding objects of one color

The classical approach for locating objects by one color from the image is rather simple and computationally fast, but unfortunately, it works well only in perfect scenarios where:

1. The fibers on the image are nearly perfectly straight, and there are no curves in the fibers.
2. There are no clusters of fibers close to each other.
3. There are no intersections between different fibers.

In order to increase the accuracy of the produced results, using our first algorithm as the basis, we have created a second algorithm that can work well and produce more accurate results in the described non-perfect scenarios. This second algorithm has the same purpose to locate and measure the length of fibers of one color in the image. It works based on skeletonization and orientation-aware traversals of the images.

This algorithm has six main stages:

1. Binary thresholding;
2. Skeletonization;
3. Locating a path that represents a fiber on a large scale;
4. Post-processing of the path produced in stage 3 by increasing the detail and making it more accurate on a small scale;
5. Marking the path on the binary image;
6. Filtering the path from the list of found fibers in case it does not match the required criteria. One of the key concepts in this algorithm is dividing the main problem to locate the fibers in stages 3 and 4 by working on a large-scale and a small-scale. This allows us to use more approximations without a significant loss of accuracy.

#### 2.4.2. Initial preparation of the images and implementation of the skeletonization process

The analyzed image is processed using binary thresholding by RGB values with an upper and a lower boundary. Next, the resulting binary image is subjected to skeletonization using our modification of the Zhang-Suen thinning algorithm (Zhang and Suen 1984). The modification was required because when the original algorithm was tested with experimental images, the produced skeletons often appeared scattered, making them unusable for further processing. The modification consists of the following: Pixels surrounded by four adjacent pixels (with a common side) with a value of true are marked as inner pixels. Then the Zhang-Suen thinning algorithm is modified to remove only pixels (set their value in the binary image to false) only if they have a common side with an inner pixel. This modification significantly reduces the number of occurrences of the problem (of the unmodified algorithm), but as a side effect, it sometimes creates lines in the skeleton with a width of two or three pixels. However, due to the way the produced skeletons are processed later, this problem was not found to create complications later in the algorithm.

The binary thresholding combined with skeletonization allows a much better definition of the region of interest (ROI) for processing than the previous method’s plain binary thresholding. The complexity of this modification of the Zhang-Suen thinning algorithm is:

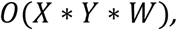

Where W is the largest thickness of an object in the image. Thickness is defined as the number of layers in an object that need to be removed to obtain the skeleton of this object.

#### 2.4.3. Graph representation and definitions

After the skeletonization process is complete, a binary image is produced where only the pixels that are part of the skeleton are marked with a value of true. This binary image is represented as a graph of type grid, and edges are built between different pixels (represented by nodes) using the same approach from the initial algorithm (finding objects by one color, classical approach). There is only one addition which is that the edges are also weighted. The following definitions will be used in this graph:

A path is defined as a finite sequence of vertices connected with a finite sequence of edges. The length (weight) of an edge in this grid is defined as the Euclidean distance between the centers of the two pixels that this edge connects. The unit for distance is the width of a pixel. The length of the path is defined as the sum of the lengths of each edge from that path. The shortest path between two different pixels(vertices) is the one with the shortest length from all possible paths between those pixels.

In the currently described algorithm, each fiber of one color from the image is represented by a path of edges from this graph. The edges from this path must have endpoints that are pixels of the produced skeletons. The length of this path will be used as the length of the found fiber. So, the goal is to locate specific paths from the grid while only traversing pixels (represented by nodes) marked as a part of the skeletons after the skeletonization process.

It is important to note that not every path from the skeletons can represent a fiber. Paths that contain loops and form closed areas on the 2D plane of the image or which have many sharp turns between adjacent edges we define as invalid to represent a fiber. To isolate those cases and produce valid paths, we use the approximation to find only paths, which are the shortest connecting a pair of endpoints, and the edges of these paths match several orientation limits (described below).

#### 2.4.4. Main traversal tool

Since this algorithm works with the shortest paths, necessary to find them in the graph of type grid. The weights of the edges are always positive (they are distances between pixels), so the Dijkstra shortest path algorithm can be used, with a complexity of (in case it is implemented effectively with a heap):

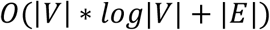

In the graphs of type grid, specific rules define the edges and their lengths. From these rules, the following information can be derived:

1. A direct edge exists between every pair of nodes if the Euclidean distance between the pixels of those nodes is smaller or equal to a defined by the user constant *J*. So, if there are nodes *V1* and *V2* connected with a direct edge and if there is a node *V3* for which the Euclidean distance between *V1* and *V3* is smaller than the Euclidean distance between *V1* and *V2*, there exist also a direct edge between *V1* and *V3*.
2. If two nodes are connected with a direct edge, the shortest path between those nodes will be that direct edge. (The edges have a meaning on the 2D plane of the image)

Using this, we have created a shortest path algorithm that is slightly faster than the Dijkstra and does not require a heap. However, it works only in the situation where edges are created by the listed rules. Our algorithm is a modification of the breadth-first search (BFS) algorithm. In this modification, the edges from a currently visited node are traversed (their endpoints put into the BFS queue) by their lengths from smallest to largest. The iteration of the nearby pixels from smallest to largest from the current pixel is done via a special construction that we have developed, which uses two nested loops and a separate queue. Since the Breadth-first search and the Dijkstra algorithms are close to each other in terms of implementation, our algorithm for finding the shortest paths in this special kind of graph can also be considered as optimization of the Dijkstra algorithm which works only in this special case. This modification of the BFS stores for each traversed pixel a reference to its so-called “parent “pixel (the pixel which ordered its traversal) in a special array. The information from this array will be used for retrieving traversal paths and orientation checks. Since the described algorithm is a modification of the Breadth-first search, it is going to have a complexity of:

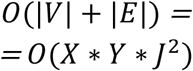

#### 2.4.5. Locating the path on a large scale

After the skeletonization process, every pixel from the image is iterated to find pixels from the skeleton that were not previously visited. When such a pixel is found, let us denote it with *P1*. Our modified version of BFS is called to find the distances from *P1* to every other pixel reachable from it. After that, the pixel with the largest distance from *P1* is marked, and let us denote this farthest pixel with *P2. P2* has interesting properties, which will be proved in the following observations. Since the produced skeletons represent fibers, formations of closed areas (loops) are extremely uncommon. Common formations of closed areas (loops) are a sign of failure in the experimental part of the research where the DNA fibers have not been stretched properly. Knowing that the traversed skeleton reassembles a tree in a certain degree and the distance between *P1* and *P2* is the longest shortest path with a starting point *P1, P2* will be an endpoint in the skeleton and in one of the farthest corners of the skeleton (Algorithm for finding a diameter in a graph of type tree). By using all these approximations, we assume that *P2* is an actual endpoint of a path that represents a fiber on the image that we need to isolate and measure its length.

After *P2* is found, with *P2* as a starting point, our version of the BFS is called again; however, in this case, this BFS is also modified with an orientation check. The orientation check consists of limiting the turning angle with a maximum value that the user can define. For this check is also used the parent array, so the turning angle to a potential for traversal next pixel can be calculated.

**Fig. 2:**
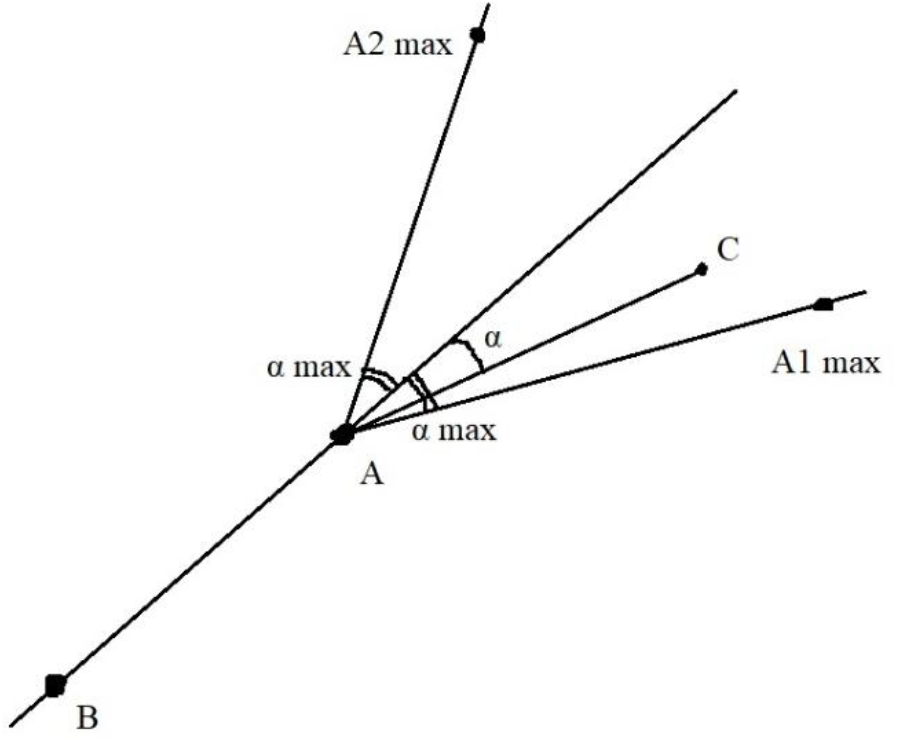
Description of the orientation check technique. *A* is the current pixel, which was the last pixel to be retrieved from the queue. *B* is the parent pixel of *A* (*B* is the pixel from which *A* was called to be pushed into the queue). *C* is the pixel which we are currently checking whether it can be reached. The current turning angle to reach *C* is represented with *α*. “*α max*” is the maximum turning degree defined by the user. *C* must lie in the area of the angle *A1AA2* in order to be traversed.

In the cases of intersecting fibers, sharp turns in the skeletons can distinguish what can be part of the same fiber, as a single fiber is extremely unlikely to be curved at a sharp angle after the stretching process.

After the run of the second version of our modified search, the longest shortest path is retrieved using the parent array, and at this stage, it represents the located fiber of one color. It is important to note that at this stage of the algorithm, we produce results that need to be accurate and missing errors only on a large scale.

#### 2.4.6. Post-processing of the produced large-scale path

After the described above method is executed, a path is retrieved whose length represents the length of the object in one color. The produced path consists mostly of edges with the maximum possible length – the defined by user “max jump distance”. That is because the previously used modification of the breadth-first search will find the endpoint of the path by using a minimum number of edges. Using the minimum number of edges to create a path, connecting fixed endpoints in the 2D plane of the image leads to using edges with the longest lengths possible, which leads to the mentioned problem. Calculating the length of skeletons of small objects in the image with a large value of the maximum jump distance creates a low-quality measurement of the length of these small objects. This can lead to problems in cases of images where it is required a large value for the maximum jump distance to be used.

So the following problem is solved by post-processing the produced path from the previous method. Let the initial path consists of *E* edges and is found with a value for the maximum jump distance defined from the user *J*. The value of *J* is set to *J/2* (rounded down to the nearest decimal value). The edges are traversed from the first to the last one. Let the current edge be *Ex*. For *Ex* is checked if the endpoint of *Ex* can be reached from the starting point of *Ex* by using the shortest path algorithm (described previously), but this time for the maximum jump distance is used the updated value of *J*. When the endpoint of *Ex* is reached, in the original path *Ex* is replaced by the path (retrieved from the last run of the shortest path algorithm of edges) connecting the endpoints of *Ex*. If the other endpoint of *Ex* is not reached, *Ex* is left in the original path. The number of times that this process is repeated can be controlled by the user by a parameter called “lower resolution bound”. This method manages to increase the resolution in *O(log2 J)* steps.

#### 2.4.7. Final work with the produced small-scale path

After the post-processing, a path containing all the pixels of a fiber from the skeleton is retrieved. It is important for the part of the skeleton that contains this path to be marked as visited, so when the next fiber in the image is searched in the initial iteration of each pixel from the image, the same fiber or fiber that overlaps parts from a previous one is not found, because this may cause an uncontrolled infinity loop. On one hand, if the entire connected component of the skeleton, which was traversed by our versions of BFS, is marked as visited, our algorithm will be limited to locate only one fiber from each connected component of a skeleton. This will make the algorithm incapable to produce accurate results in cases of clusters and intersections, which was defined as a main goal of this algorithm. On the other hand, if only the pixels from the found path after the post-processing are marked as visited, the list of found fibers will be contaminated by many objects that are nearly identical due to the fact that the skeletons are often not with a width of a single pixel. For this problem, we have implemented a simple solution that consist of marking as visited every pixel in a circle around every pixel from the path of the located fiber. The radius of these circles currently is a constant named as “*Cleaning Offset*” and its value is defined by the user. In case this solution needs to be further improved a value from a distance map acquired from a distance transform can be used instead of this constant defined by the user.

Finally, the length of the path is calculated and this length is used as the length of the fiber. Also the found fibers can be filtered by saving the information only about those fibers from the image that have a length not smaller than a defined by the user minimum value. This is important, because in the images exist many small artefacts that can contaminate the produced results. In this algorithm can be included in the future many other different options to filter the produced paths. At the moment this length filter manages to remove artefacts in most cases.

#### 2.4.8. Complexity of the approach with skeletonization for locating objects of one color

In the best-case where there are no clusters or intersections of objects and for each connected component is found only one fiber, the complexity of this entire algorithm will be:

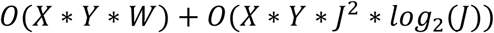

However, often there may be found more than one fiber in a single connected component. This leads to traversing pixel from a skeleton more than once, which we have assumed as impossible in the best case. So in the worst case the complexity of this second algorithm for locating objects of one color will be:

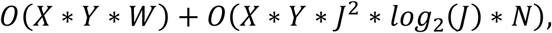

where *N* is the number of fibers that were found in the image before their filtering by length and other possible parameters in the future.

### 2.5. Finding objects consisting of two colors

The method allows to find objects of interest containing segments of two different colors. This method requires, references to settings profiles for finding objects of each color. Since the image may contain artefacts between the segments of different color a parameter “maximum merge distance” is introduced so these segments can be merged if the distance between them is smaller than that value.

The distance between two single-color objects can be calculated in two different ways, depending on their types. If the objects were found using the simpler classical approach, then the distance is presented as the distance between the parallel segments forming the rectangles that represent the single-color objects. If the objects were found using the skeletonization approach the distance between the two objects is presented as the smallest distance between the endpoints of the paths of these two objects.

Initially, single-color objects are found and stored into two different sets using the algorithms described above. Then they need to be connected into pairs so each pair has an object of the first and an object of the second color and the distance between these two objects must be smaller than the value defined by the user. These pairs represent the more complex two-color objects. Each single-color object can belong only to one pair. Not only a maximum number of pairs must be found, but the objects of different colors must be matched in the best possible way. The best possible way is defined when the sum of the distances between the matched objects is minimal. So, the problem is reduced to the minimum cost matching problem.

A bipartite graph is being built with edges containing capacity and cost to be run a unit of flow through them. There are two special nodes created – source and a sink. Every object of both colors represents a node. Let *N* be the number of objects from both colors that need to be paired. Then:

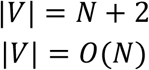

Edges from the source to every object of the first color and from every object from the second color to the sink are built with a capacity of 1 and a cost to run flow through them of 0. Between objects of two different colors an edge exists if the distance between those objects is smaller than the defined by the user maximum, the capacity of those edges are set again to 1 and the cost to run a flow through them is that distance. Also, for every one of those edges an opposite edge is built with a capacity of 0 and a cost of *-c*, where *c* is the cost of the original edge. To store this graph is used an adjacency list. We are creating edges for a possible combination between two objects, then:

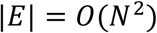

After the bipartite graph is built the minimum cost maximum flow is found. The Successive Shortest Path algorithm is being used to calculate the minimum cost maximum flow. For each run of the Successive Shortest Path Algorithm is chosen the Shortest Path Faster Algorithm (an improvement of the Bellman–Ford algorithm) (Duan 1994). During the execution of the algorithm, the edges through which flow is being run, are being labeled so the resulting pairs of objects of different colors can be obtained. The Successive Shortest Path algorithm can be seen as an implementation of the Ford-Fulkenson algorithm for maximum flow (Ford and Fulkerson 1956), where for finding each path is used a shortest path algorithm (in our case - Shortest Path Faster Algorithm). The worst-case complexity of the SPFA algorithm is:

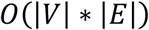

So, in the worst case this matching algorithm will perform in:

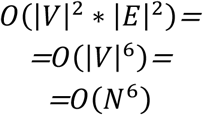

### 2.6. DNA fiber labeling

DNA fiber analysis was performed as described by before (Schwab and Niedzwiedz 2011, Hristova, Stoynov et al. 2020). Briefly, exponentially growing PC3 cells were first incubated with 25 µM chlorodeoxyuridine (CldU) for 10 minutes and then with 250 µM iododeoxyuridine (IdU) for 30 minutes. Hydroxyurea (HU) at the indicated concentrations was added simultaneously with the secon label (IdU). Spreads were prepared from 2500 cells (suspended in PBS at 1×10^6^ cells/ml). Cell lysis was carried out in fiber lysis solution (50 mM EDTA and 0.5% SDS in 200 mM Tris-HCl, pH 7.5). DNA fibers were spread by tilting the slides ∼ 25 degrees until the drop of the fiber solution reached the bottom of the slide and was let to dry. Dried slides were processed immediately. Slides were suspended in 2.5 M HCl for 80 min, washed in PBS, and then incubated in blocking buffer (5% bovine serum albumin in PBS) for 40 min. Primary antibodies - mouse anti-BrdU antibody (Becton Dickinson, cat # 347580) to detect IdU and rat anti-BrdU antibody (Abcam cat# Ab6326) to detect CldU - were diluted in blocking buffer and applied overnight. Slides were washed several times in PBS, incubated with secondary antibodies for 60 min and mounted with ProLong Gold anti-fade reagent (Molecular Probes). Images were acquired with Axiovert 200M microscope (Carl Zeiss) equipped with Axiocam MR3 camera (Carl Zeiss).

## 3. Results and discussion

### 3.1. Platform architecture

The platform “DNA size finder” was developed as a desktop application in Java. The application uses a simplified version of the layered architecture with 3 distinct layers:

1. Graphical user interface (GUI) – The entire visual interface with the user.
2. Domain – The core part of the program where all image processing objects are implemented (loaded images, object groups, settings). The domain layer is created by following the factory object-oriented pattern.
3. Common – In this layer are implemented the resources, that classes from the other layers can use, such as enumerations and our own libraries with mathematical functions, custom containers and etc.

### 3.2. Graphical user interface

The graphical user interface was implemented by using the standard Java Swing API. The Swing API was chosen, because it is an easy way to provide a stable cross-platform GUI in a Java application without using any third-party libraries. The graphical user interface is organized in a main window of the application from which different dialog windows can be called. The main window is organized into 5 different parts:

1. Main panel – The largest central part of the window, where the images are visualized.
2. Workspace explorer – A hierarchical menu where all instances (loaded images, found object groups and settings) that are being used by the user are ordered.
3. Console – A small panel located below the workspace explorer where detailed info is displayed.
4. Horizontal toolbar – Located at the top part of the window, it contains buttons that display drop-down menus to access features associated to the entire platform. (the factory mode).
5. Vertical toolbar – Located at the left part of the window, in it tools are located to manipulate the results produced by the program.

**Figure 3.**
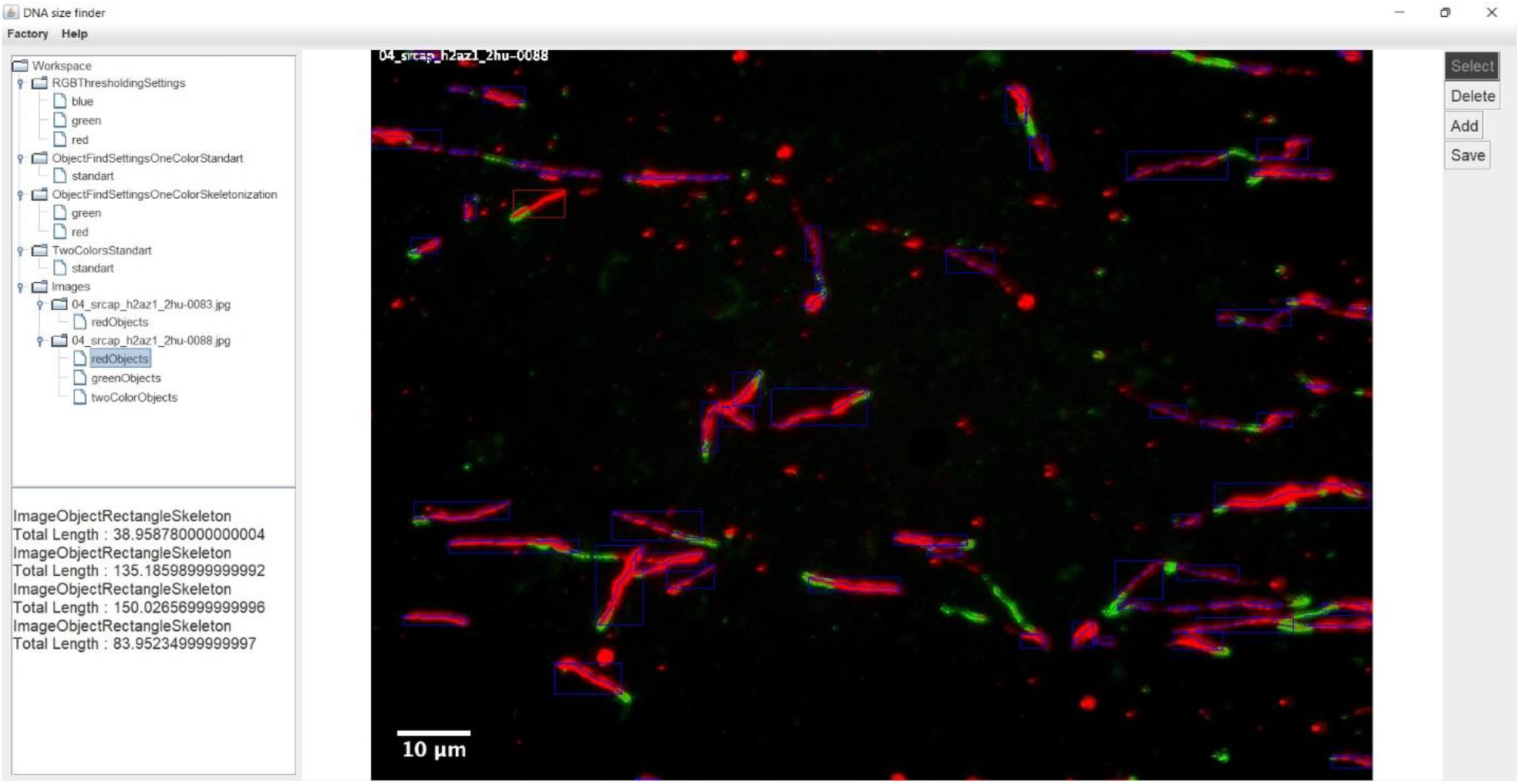
An image of the graphical user interface of the platform “DNA size finder”.

### 3.3. The settings system

The platform “DNA size finder” uses a hierarchical settings system to store the parameters used by the algorithms for the analysis of the microscopic images. Each algorithm for image analysis has its own settings type in our platform. The settings types are divided into three different groups (classes):

1. “Simple settings” – These settings cannot be used to locate objects from the images. Their main purpose is to be used by advanced settings types. At the moment the RGB thresholding is represented by settings of this type.
2. “Single object find settings” – These settings are needed to find objects in the image by using algorithms implemented by simple settings. Examples for such settings are the ones for finding single-color objects via the classical or skeletonization approaches.
3. “Multiple object find settings” – These settings affect the more complex types of algorithms, which in turn, use the algorithms modified by the simple and single object find settings types. An example from this class of settings is the settings for locating fibers with two colors.

For every setting type many profiles (instances) can be created by the user with different parameter values, so when a user wants to analyze an image, a pre-created settings profile can be chosen. Each settings profile is automatically saved in the permanent memory of the computer and loaded with the start of the program. The settings instances can be easily transferred as a file from one computer to another.

### 3.4. The factory mode

The factory mode is an experimental feature in our platform. It allows the researchers by using an already defined instance of an object find settings to analyze a large database of many images completely automatically. This mode allows entire databases to be analyzed without a researcher supervision.

### 3.5. Testing the application

We tested DNA size finder on a set of images obtained in control cells and ones treated with the low doses of the replication inhibitor HU during the incubation with the second label. Cells were subjected to DNA fiber labeling and the images were analysed either manually or using the DNA size finder. Data indicated that the application produced similar results to manual counting both in “normal” (user corrected) mode and in the fully automated “factory” mode (Fig. 4). We observe that the application produced more extreme measurements, which seem to be removed by the trained expert, not necessarily in an unbiased way.

**Fig. 4.**
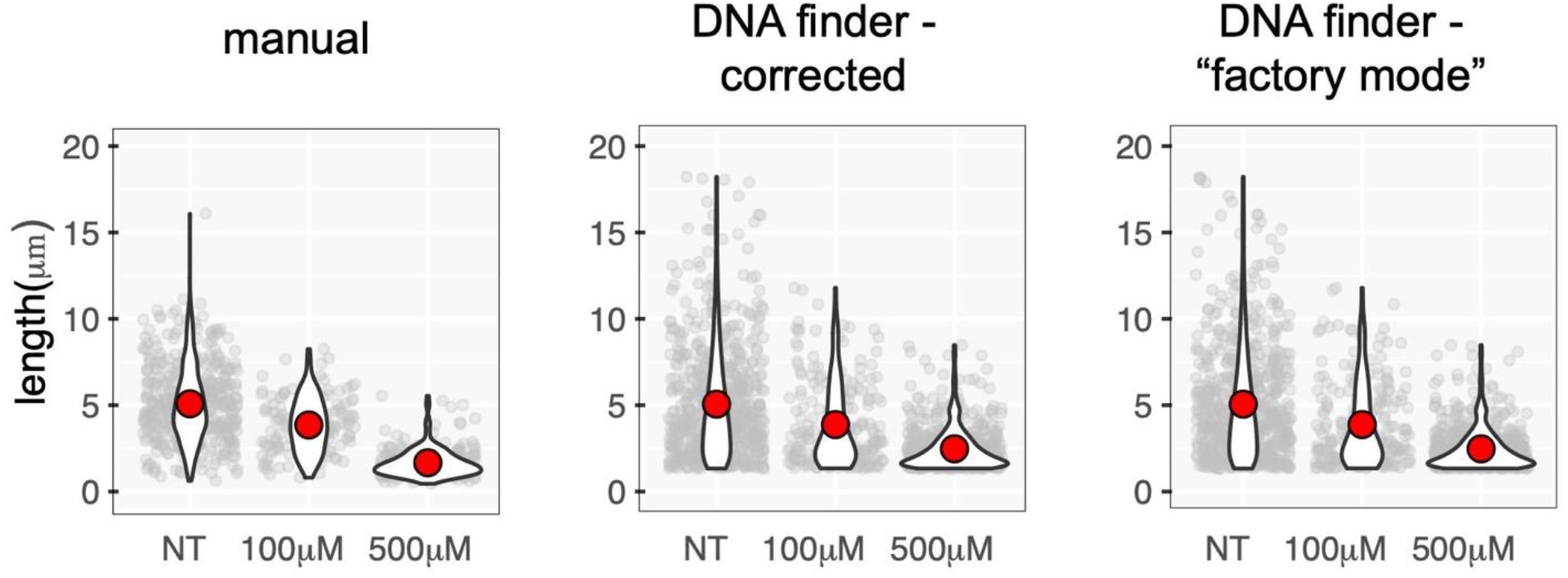
Analysis of DNA fibers (length of second label) obtained from cells treated with the indicated concentration of HU, subjected to manual analysis, or analysis by the DNA size finder in user supervised and fully automated modes. The red dot on violin plots indicate the mean of all measurements (plotted in grey behind the violin plots).

## 3.6. Discussion

The need to study replication dynamics and the difficulty of the analysis of images obtained via DNA fiber labeling has spurred the interest in creating automated solutions to the problems in the last two years (Ghesquiere, Elsherbiny et al. 2019, Mohsin, Arnovitz et al. 2020). Here, we describe the algorithm and provide as an example a lightweight Java application to automatically analyze single DNA molecule images. DNA size finder significantly simplified the analysis of the experimental data while increasing reliability by the standardized measurement of a greater number of DNA molecules. Unlike other similar solutions it does not rely on other software tools, (except freely available Java Runtime environment), it is multi-platform, does not need a complicated set up and uses explicitly described algorithms. DNA size finder offers several modes of operations – single or dual color evaluation, user corrected or fully automated image analysis.

In addition, by adjusting the object detection settings DNA size finder operation can be user optimized to the specific experiment. Finally, the approaches applied may be helpful in other settings requiring image analysis.

## 4. Data Availability

The application is free to use and can be accessed at the following link: https://github.com/Teodor03/DNA-size-finder

